# Stapled peptides based on Human Angiotensin-Converting Enzyme 2 (ACE2) potently inhibit SARS-CoV-2 infection *in vitro*

**DOI:** 10.1101/2020.08.25.266437

**Authors:** Francesca Curreli, Sofia M B Victor, Shahad Ahmed, Aleksandra Drelich, Xiaohe Tong, Chien-Te K Tseng, Christopher D. Hillyer, Asim K Debnath

## Abstract

SARS-CoV-2 uses human angiotensin-converting enzyme 2 (ACE2) as the primary receptor to enter host cells and initiate the infection. The critical binding region of ACE2 is a ∼30 aa long helix. Here we report the design of four stapled peptides based on the ACE2 helix, which is expected to bind to SARS-CoV-2 and prevent the binding of the virus to the ACE2 receptor and disrupt the infection. All stapled peptides showed high helical contents (50-94% helicity). On the contrary, the linear control peptide NYBSP-C showed no helicity (19%). We have evaluated the peptides in a pseudovirus based single-cycle assay in HT1080/ACE2 and human lung cells A549/ACE2, overexpressing ACE2. Three of the four stapled peptides showed potent antiviral activity in HT1080/ACE2 (IC_50_: 1.9 – 4.1 µM) and A549/ACE2 cells (IC_50_: 2.2 – 2.8 µM). The linear peptides NYBSP-C and the double-stapled peptide StRIP16, used as controls, showed no antiviral activity. Most significantly, none of the stapled peptides show any cytotoxicity at the highest dose tested. We also evaluated the antiviral activity of the peptides by infecting Vero E6 cells with the replication-competent authentic SARS-CoV-2 (US_WA-1/2020). NYBSP-1 was the most efficient preventing the complete formation of cytopathic effects (CPEs) at an IC_100_ 17.2 µM. NYBSP-2 and NYBSP-4 also prevented the formation of the virus-induced CPE with an IC_100_ of about 33 µM. We determined the proteolytic stability of one of the most active stapled peptides, NYBSP-4, in human plasma, which showed a half-life (T_1/2_) of >289 min.

## Introduction

A novel and highly pathogenic coronavirus named SARS-CoV-2 was first identified in Wuhan, China, in 2019. It causes Coronavirus disease 2019 (COVID-19), and the resultant rapid global spread is a substantial threat to public health and the world economy. On March 11, 2020, the World Health Organization (WHO) declared the COVID-19 outbreak to be a global pandemic. As of November 2, 2020, 216 countries were affected by this disease (https://www.worldometers.info/coronavirus/countries-where-coronavirus-has-spread/), with more than 47 million positive cases, including more than 1.2 million deaths. In the US, there are more than 9.2 million confirmed cases with more than 231,000 deaths (https://coronavirus.jhu.edu/data). At present, no vaccines or therapeutics are available to prevent or treat COVID-19. However, Remdesivir and some corticosteroids, such as dexamethasone, are providing some benefit to COVID-19 patients. Several vaccines and other repurposed therapeutics are also in clinical trials. SARS-CoV-2, like its predecessor SARS-CoV, uses its receptor-binding domain (RBD) in the S1 spike protein to initiate entry by binding to human host receptor angiotensin-converting enzyme 2 (ACE2). Recently, soluble human ACE2 (hACE2) has been proposed as a competitive interceptor of SARS-CoV^1^, and a recombinant hACE2 (rhACE2) is now undergoing a clinical trial^2^. However, a recent report indicated that in both humans and mice, rACE2 had a fast clearance rate, and the half-life was only hours by pharmacokinetic studies^3, 4^. Recently, it was shown that when ACE2 was fused with an Fc fragment ACE2-Ig it showed high-affinity binding to the RBD of both SARS-CoV and SARS-CoV-2. ACE2-Ig also exhibited potent neutralization of SARS-CoV-2 with improved pharmacological properties in mice^5^. Despite the potential of recombinant ACE2, in a dire global pandemic situation, protein-based therapy may not be suitable for use, especially in more developing nations, primarily due to the high cost and stability of protein-based therapy. Therefore, we sought to design therapeutics that are more stable and less expensive to make for wider distribution. Recently, Zhang et al. reported a 23-mer peptide (SBP1) from the ACE2 α1 helix sequence, which showed a potent binding affinity (K_D_=47 nM) to the SARS-CoV-2 RBD. However, there was no antiviral activity data reported^6^. We envision that this type of short linear peptides will not be stable against proteolysis and, most significantly, may not have the right secondary structure required for efficient binding with the SARS-CoV-2 RBD.

Recent Cryo-EM and X-ray structures of the SARS-CoV-2 RBD with full-length ACE2 revealed structural details of the binding interactions of these two proteins^7-10^. A close inspection revealed that the interaction surface is long and flat, which may not be conducive to design small-molecule drugs to prevent protein-protein interactions. Interestingly, however, the structure also revealed that one of the long α-helix (herein termed Helix-1) interacts with the RBD consisting of multiple binding “hot spots.” Based on this structural insight, we designed stapled peptides based on the binding site of SARS-CoV-2 RBD on the ACE2 receptor (Helix-1), which are expected to act as a decoy receptor preventing the binding of SARS-CoV-2 to the host receptor ACE2 (**Figure 1**) hence virus entry to host cells. Targeting the entry pathway of SARS-CoV-2 is ideal for both prevention and treatment as it blocks the first step of the viral life cycle.

**Figure 1.**
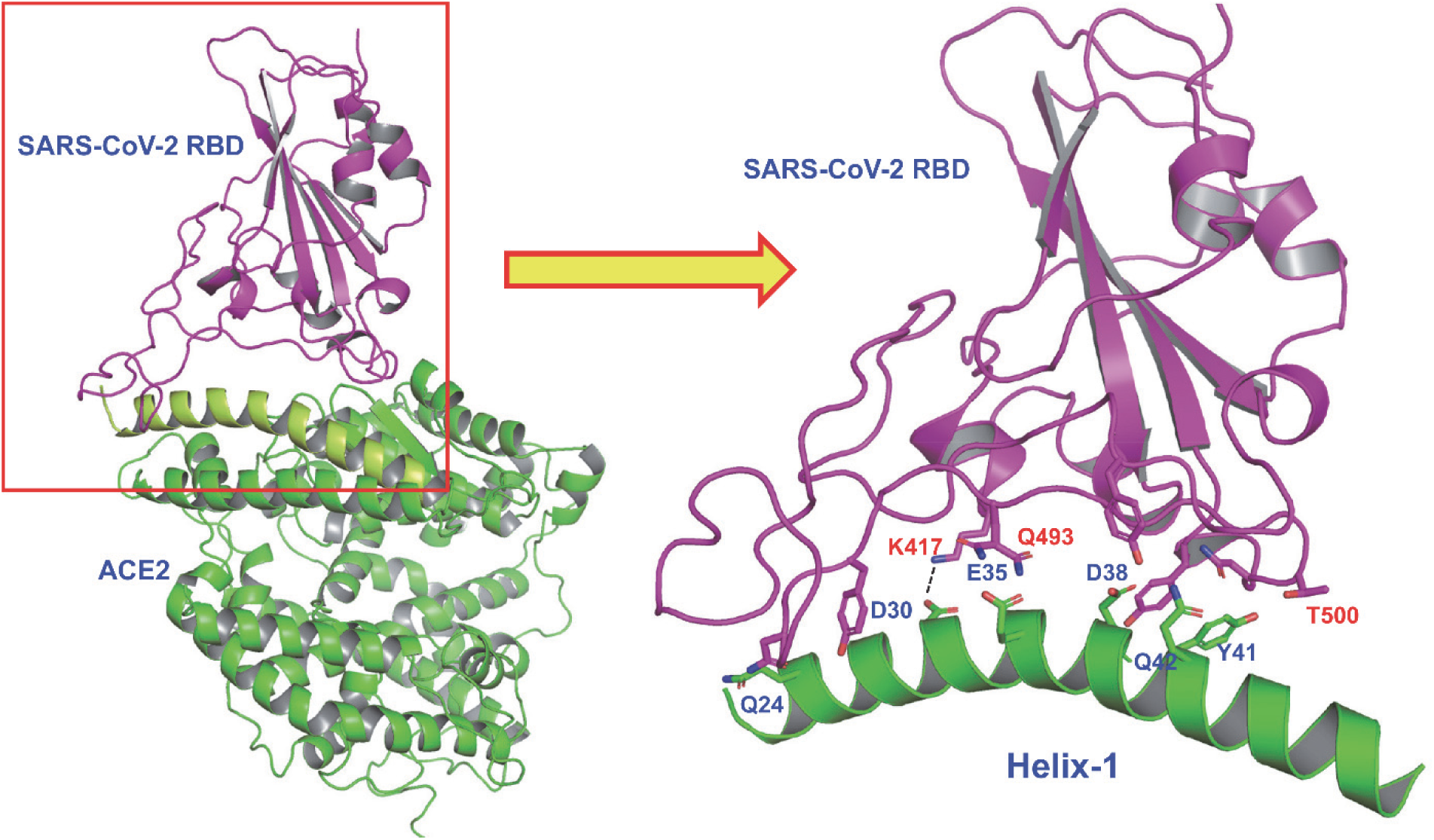
X-ray structure showing binding of SARS-CoV-2 RBD (violet) with ACE2 receptor (green). The detailed interactions of Helix-1 of ACE2 with SARS-CoV-2 RBD are indicated in the figure on the right.

Protein-protein interactions (PPI) between host and virus are valuable targets to inhibit viral replication. However, the large and flat interacting surfaces of PPI often preclude the use of small molecules as drugs to disrupt these PPI. Larger biologics, such as peptidomimetics (e.g., hydrocarbon-stapled-peptide mimetics), are, on the other hand, promising inhibitors of PPI that were previously intractable^11-14^. Stapled peptides are typically derived from the α-helix from the binding interface, and they are locked into bioactive conformations through the use of chemical linkers. Stapling enhances α-helicity of unstructured short peptides in solution, improves resistance against proteolytic digestion, as well as potency, and often improves cell penetration^15-21^. One of these stapled peptides, ALRN-6924, is in various clinical trials for the treatment of lymphomas (Phase 1b/2), peripheral T-cell lymphoma (Phase 2a), acute myeloid lymphoma (AML, Phase 1), and advanced myelodysplastic syndrome (MDS, Phase 1)^22^

Here, we took advantage of Helix-1 of ACE2 to design peptide-based therapeutics for COVID-19. However, Helix-1 is expected to lose its helical conformation in solution; therefore, it will also lose the orientation of most, if not all, of the binding site residues to make any proper interaction. To counter this, we adopted the very well-studied concept of hydrocarbon stapling techniques mentioned above to reinforce the bioactive helical conformation of Helix-1 (**Figure 2**). We hypothesize that stapled peptides will be α-helical and will be proteolytically stable. They are expected to bind to the SARS-CoV-2 RBD and inhibit virus entry to cells.

**Figure 2.**
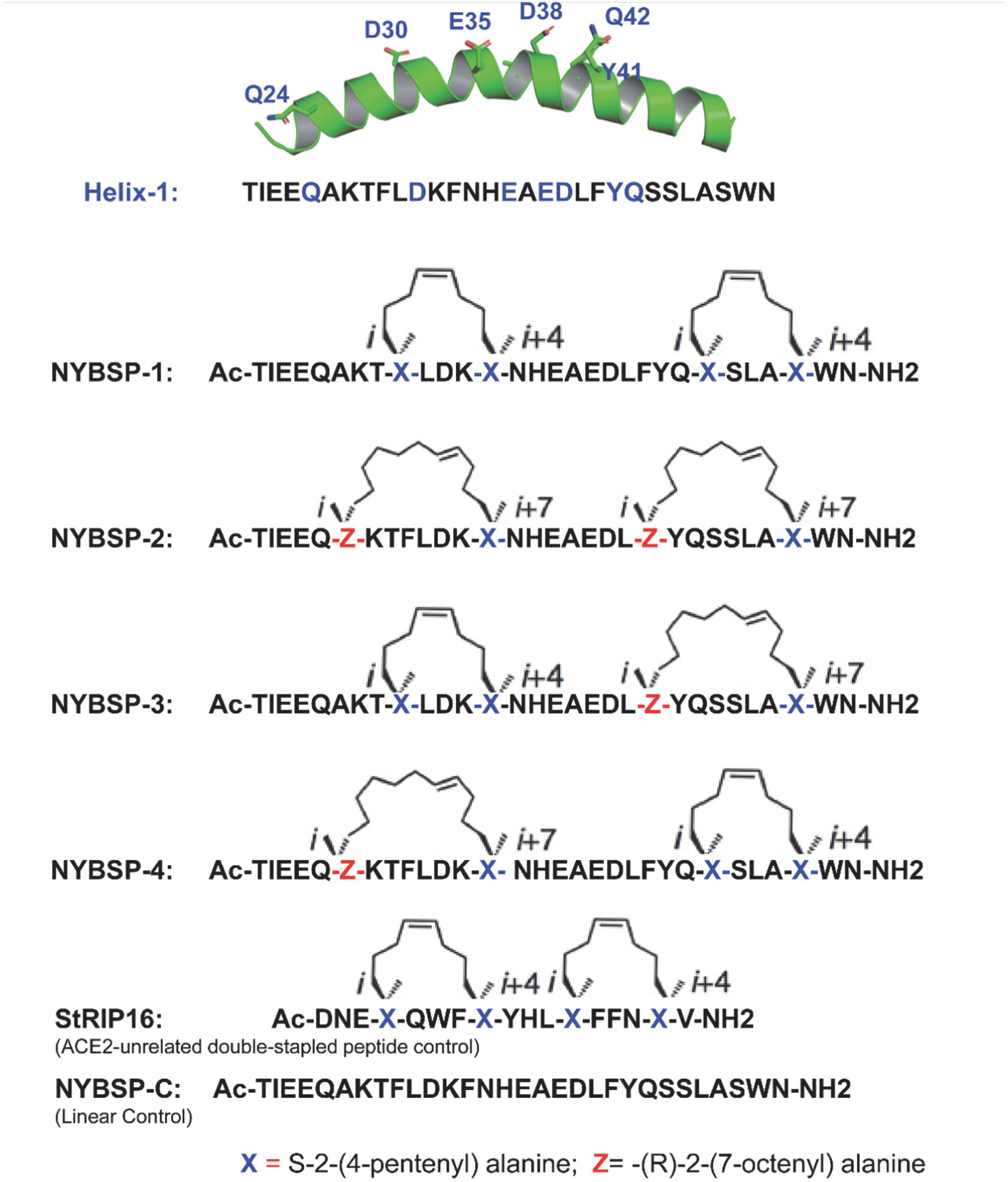
Design of Double-stapled peptides as anti-SARS-CoV-2 inhibitors. The linear peptide, NYBSP-C, and an ACE2-unrelated double-stapled peptide, StRIP16, ^29^ were used as controls.

In this report, we present the design of four such stapled peptides based on Helix-1 of ACE2. Three of these peptides showed potent SARS-CoV-2 inhibitory activity in an *in vitro* pseudovirus and in an authentic virus infectivity assay. These stapled peptides have the potential as a therapeutic for COVID-19 by functioning as a decoy of ACE2 binding site for SARS-CoV-2 and preventing the virus from binding to the ACE2 receptor; thereby preventing virus entry and infection.

## Results & Discussions

### Design of double-stapled peptides

Recently solved structures by Cryo-EM and X-rays provided intricate details on how SARS-CoV-2 RBD binds to the host receptor ACE2. The major binding sites are located on Helix-1 of ACE2 (**Figure 1**). A recent report indicated that a computationally designed short peptide from Helix-1 of ACE2 showed a nanomolar binding affinity to SARS-CoV-2 RBD^6^. However, the usefulness of such a linear peptide in isolation as a drug is yet to be proven. In general, when a helix is taken out as a peptide from a protein, it loses its secondary structure. These peptides are also prone to proteolytic degradation. On the contrary, if such small peptides are constrained by hydrocarbon stapling, an enhancement of helical content can be achieved together with resistance to proteolysis and improved binding affinity and biological activity^11-21, 23, 24^. Therefore, we envisioned that the stapling of Helix-1 will render it α-helical, resistant to proteolysis, and, therefore, a highly potent inhibitor of SARS-CoV-2.

### i/i+4 and i/i+7 hydrocarbon stapling

We used the well-established stapling techniques to synthesize hydrocarbon-stapled peptides^25-27^ from Helix-1 of ACE2, which are expected to maintain high α-helicity, have high binding affinity to SARS-CoV-2 RBD, and have improved antiviral activity and stability from proteolysis. We decided to use double stapling because of the relatively longer helical region that binds to the SARS-CoV-2 RBD. Double stapling has been reported to confer striking protease resistance^20, 28, 29^. Furthermore, Mourtada et al. recently reported 2-4 fold enhancement of antimicrobial potency by double-stapled peptides over single stapling^23^. Double stapling was also reported to achieve improved pharmacokinetic profiles, including oral absorption of peptide-based HIV-1 fusion inhibitors^28^. For the *i,i+4*, and *i,i+7* stapling, we used non-natural amino acids as illustrated in **Figure 2** to introduce the hydrocarbon staples to Helix-1. The stapling sites were selected based on the binding site information of SARS-CoV-2 RBD with ACE2 (Helix-1) from the Cryo-EM and X-ray structure mentioned before so that they do not interfere with the critical binding site residues. We have synthesized (CPC Scientific, Inc.) four stapled peptides, as depicted in **Figure 2**. We also synthesized the linear peptide, NYBSP-C, as a control. Besides, we purchased StRIP16 ^29^, which is a Rab8a GTPase-binding double-stapled peptide and unrelated to ACE2, to use as a control.

### Effect of double-stapling on the helical propensity

Due to the extended size of Helix-1, we decided to double staple Helix-1 of ACE2 to preserve the α-helical content of the peptides. To determine the helical propensity of these peptides, we resorted to CD spectroscopy for analyzing secondary structures. The Data in **Figure 3(A)** indicate that all the stapled peptides had the maxima at 190 nm and two minima at 208 and 222 nm, confirming that all these stapled peptides had α-helical structure. The % helicity calculation shows remarkable improvement over the linear peptide. The data in **Figure 3(B)** indicate that NYBSP-1 had the highest helical propensity of 94%. Stapled peptides NYBSP-2 and NYBSP-4 had % helicity of 61 and 80, respectively. NYBSP-3 showed the lowest % helicity (50%) among the four stapled peptides. The control peptide, NYBSP-C, as expected, showed no indication of α-helical propensity, with only 19% helicity. The control stapled peptide, StRIP16, was reported to have 38% helicity.^29^

**Figure 3.**
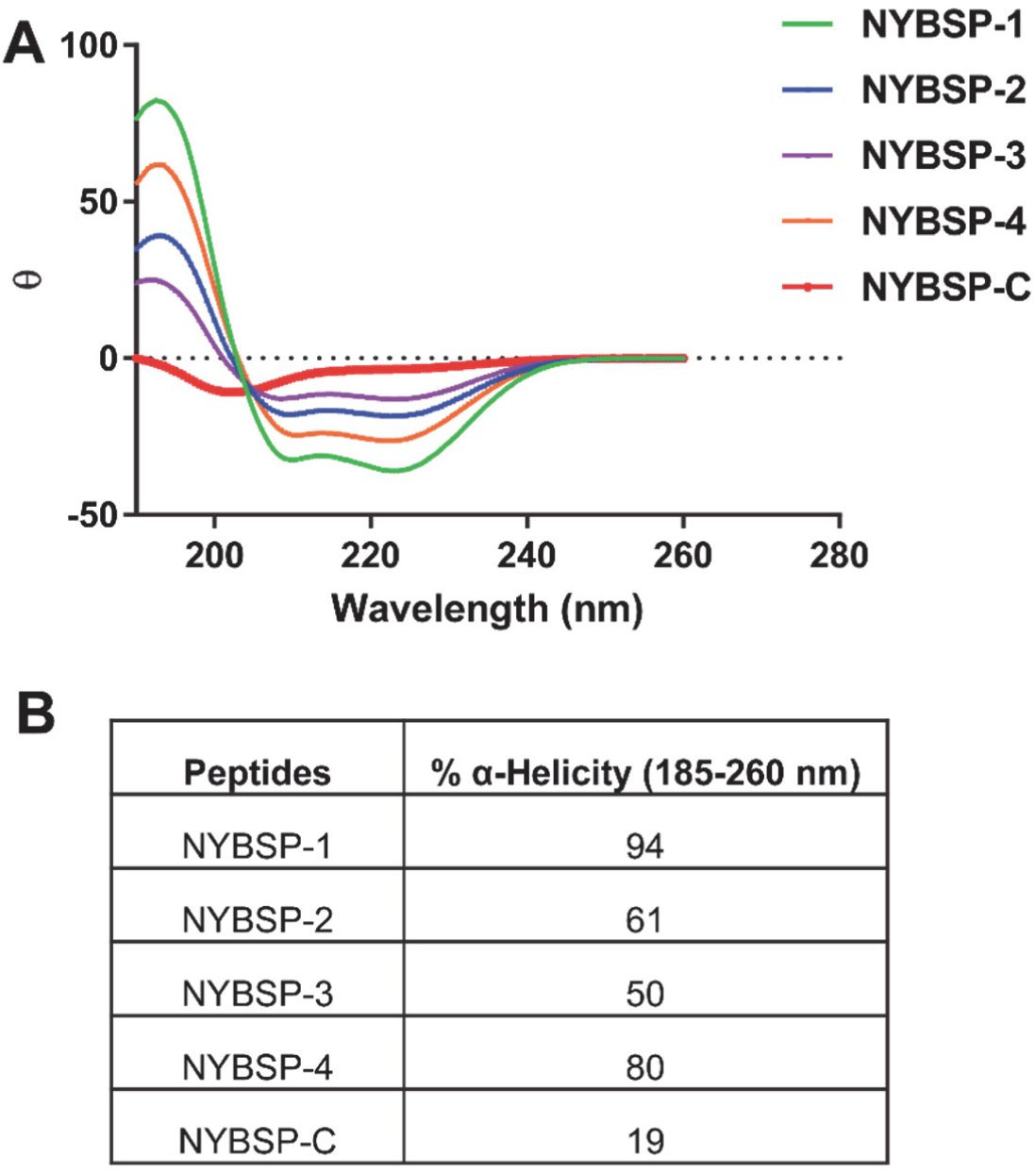
α-Helical propensity measured by CD spectroscopy **(A)** CD spectra, **(B)** % Helicity of the double-stapled peptides, and the linear control peptide.

### Binding affinity of a double-stapled peptide by SPR analysis

We used SPR to determine the binding affinity of one double-stapled peptide, NYBSP-4, with SARS-CoV-2 RBD. This method is useful in measuring the binding constant (K_D_) as well as *k*_on_ (also known as association constant, *k*a) and *k*_off_ (also known as dissociation constant, *k*_d_). We also used the linear peptide, NYBSP-C, as a control. Various concentrations of biotinylated SARS-CoV-2 RBD was manually printed onto the Chip (immobilization) via biotin-avidin conjugation. Peptides were passed through the chip surface, and the signal changes (in AU) of each peptide at varied concentrations were recorded (**Figure 4A and B)**. The resulting data were fit to a 1:1 binding model, and the binding affinity K_D_ and kinetic parameters *k*_on_ and *k*_off_ of the interaction of SARS-CoV-2 RBD with NYBSP-C and NYBSP-4 were determined (**Figure 4C**). The K_D_ value of NYBSP-4 was 2.2 µM, which was very much in line with the antiviral activity we observed against SARS-CoV-2 pseudovirus (**Table-1**). As expected, the linear peptide, NYBSP-C, used as a control showed poor binding (73.7 µM), which was about 33.5-fold lower than NYBSP-4. The SPR analysis also confirmed that the double-stapled peptides bind directly to the SARS-CoV-2 RBD.

**Figure 4.**
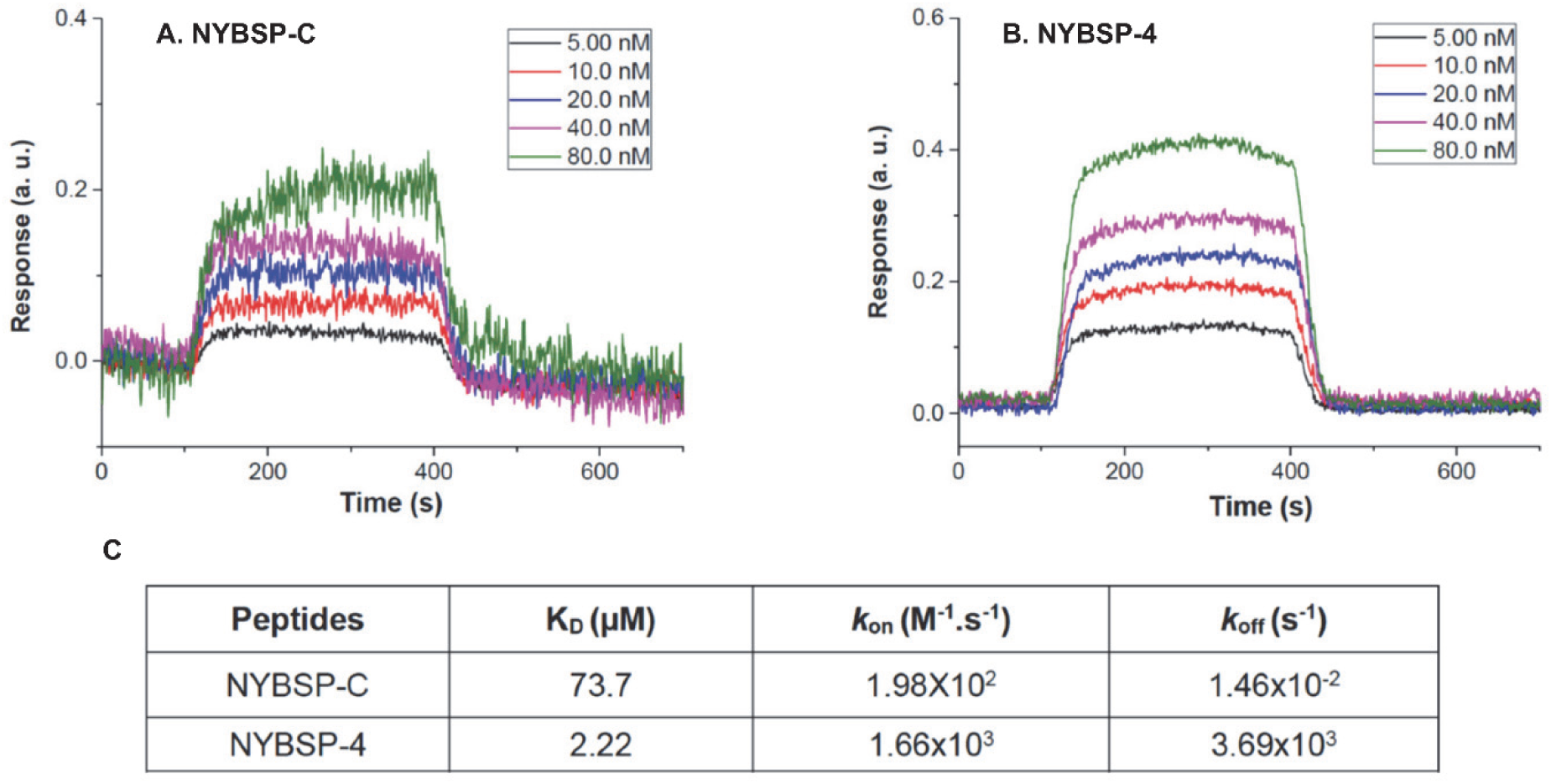
Evaluation of binding affinity of peptides to SARS-CoV-2 RBD by SPR. Kinetics fitting curve (sensogram) of SARS-CoV-2 RBD to (A) NYBSP-C, a linear control peptide; (B) NYBSP-4, a double-stapled peptide; and (C) the binding affinity K_D_ and kinetic parameters *k*_on_ and *k*_off_ of NYBSP-C and NYBSP-4.

### Proteolytic stability of stapled peptides in human plasma

Proteolytic stability remains a potential concern in developing peptide-based therapy. We wanted to address this problem by introducing double stapling in Helix-1 of the ACE2 receptor. Double stapling is known to enhance protection from proteolysis. However, it was prudent for us to test how the stapled peptides that we designed withstand the proteolysis in human plasma. The peptides were incubated with human plasma at 37°C, and the fragments formed from proteolysis were measured by LC-MS/MS at different time points (0,10, 30, 60, and 120 min). We used a small molecule drug, propantheline bromide, as an assay control. The data in **Figure 5** shows that stapled peptide NYBSP-4 is highly stable during the time used for the assay. The calculated half-life (T_1/2_) of NYBSP-4 was >289 min, but surprisingly, the linear control peptide NYBSP-C also showed similar stability (T_1/2_ >289 min). Usually, small peptides are known to get proteolytically cleaved quickly. The unusual stability of the linear peptide in human plasma in our test is yet to be understood.

**Figure 5.**
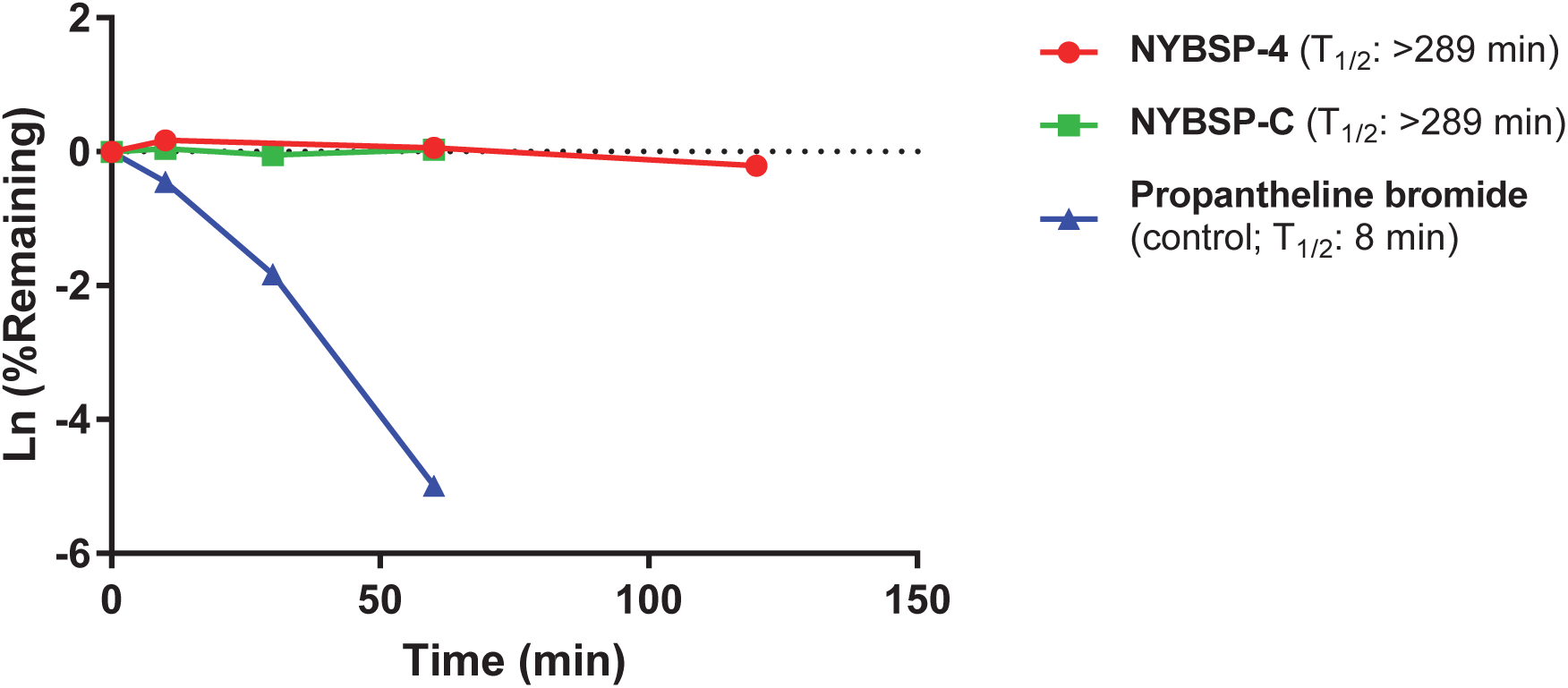
The plasm stability of double-stapled peptide NYBSP-4 (red) and control linear peptide, NYBSP-C (green). Propantheline bromide (a small molecule antimuscarinic agent) was used as an assay control.

### Validation of the SARS-CoV-2 pseudovirus

We prepared SARS-CoV-2 pseudovirus capable of single-cycle infection by transfecting HEK293T/17 cells with an HIV-1 Env-deleted proviral backbone plasmid pNL4-3ΔEnv-NanoLuc and pSARS-CoV-2-S_Δ19_ plasmid^30^. The incorporation of the spike protein in the pseudovirus was validated by Western blot analysis using the SARS Spike Protein Antibody (Novus Biologicals) (**Figure 6A**). This antibody targets explicitly amino acids 1124-1140 of the spike protein S2. We found a specific band at 80 kDa, which identifies the subunit S2 and a second band at about 190 kDa, which corresponds to the full-length S protein (S1 + S2), as previously reported ^31, 32^, confirming the incorporation of the SARS-CoV-2 S protein in the pseudovirus. Moreover, the correlation of SARS-CoV-2 pseudovirus with the expression of the ACE2 receptor was analyzed by infecting five different cell types including the human fibrosarcoma HT1080 cells, overexpressing the ACE2 (HT1080/ACE2) and the parental HT1080 cell type, the human lung carcinoma cells A549 overexpressing ACE2 (A549/ACE2) and the parental A549 cell type, and the HeLa cells which do not express the hACE2. The cells were infected with the same volumes of the supernatant-containing pseudovirus SARS-CoV-2. As expected, we found no infection of the HeLa cells (**Figure 6B**), while we detected a low level of infection of the HT1080 and A549 cells (1.5×10^5^ and 9.2×10^4^ RLU, respectively). The HT1080/ACE2 supported high levels of SARS-CoV-2 infection. In fact, we detected about 10^8^ RLU, which corresponds to a 665-fold higher infection than what we detected for the parental cell type HT1080. The infection detected for the A549/ACE2 cells was moderate (about 8×10^5^ RLU) compared to HT1080/ACE2 but about 9-fold higher than what we detected for the parental cell type A549. These results were confirmed by analyzing the expression levels of the ACE2 receptor in the different cell lines by western blot (**Figure 6C**). As shown in Blot-1, we found that ACE2 expression was undetectable in the parental cell lines HT1080, while it was overexpressed in HT1080/ACE2 cells. The lower infection detected in A549/ACE2 cells suggested a lower ACE2 expression in these cells. For this reason, in Blot-2, we loaded a higher amount of proteins (75 µg). As expected, the expression levels of ACE2 were undetectable in the parental cell lines A549 and the HeLa cells. The expression was moderate in the A549/ACE2 cells compared to HT1080/ACE2. These data confirmed that SARS-CoV-2 pseudovirus infects the cells through its interaction with the ACE2 receptor.

**Figure 6.**
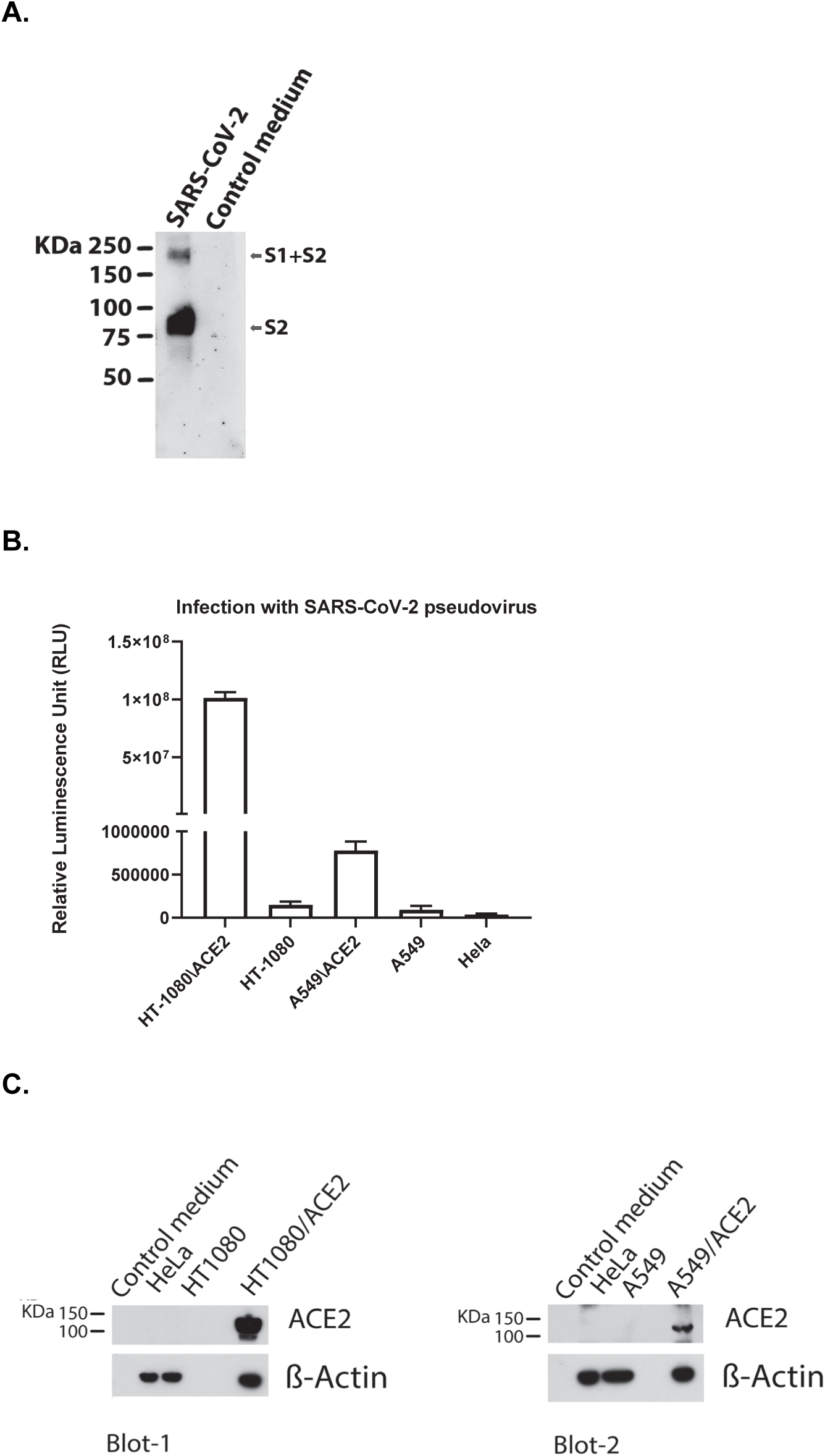
Validation of SARS-CoV-2 pseudovirus and ACE2 expression in different cells. (A) Western blot analysis to validate the incorporation of the S spike protein in the SARS-CoV-2 pseudovirus using the SARS Spike Protein Antibody (Novus Biologicals), which targets the spike protein S2. (B) Infection of cells expressing different levels of the ACE2, with the same amounts of SARS-CoV-2 pseudovirus. Columns represent the means ± standard deviations (n =3). (C) Immunoblot analysis of cell lysates to evaluate ACE2 expression. β-Actin was used as a loading control.

### Anti-SARS-CoV-2 activity and cytotoxicity of double-stapled peptides in cell-based assays

We evaluated the anti-SARS-CoV-2 activity of the double-stapled peptides by infecting the two cell types overexpressing the hACE2, the human fibrosarcoma HT1080/ACE2 cells and the human lung carcinoma cells A549/ACE2 cells, with a SARS-CoV-2 pseudovirus that was pre-treated with escalating concentrations of the peptides for 30 minutes. The linear peptide NYBSP-C, an unrelated double-stapled peptide StRIP16 ^29^, and an anti-ACE2 mAb (AC384 from Adipogen Life Sciences) were used as controls. The concentration required to inhibit 50% (IC_50_) of SARS-CoV-2 pseudovirus infection was calculated. The results are shown in **Table 1**. The dose-response plots (**Figure 7**) indicate that all the double-stapled peptides efficiently inhibited the infection of HT1080/ACE2 cells and A549/ACE2 cells by the SARS-CoV-2 pseudovirus with low μM potency. Three of the four stapled peptides displayed potent antiviral activity in both cell lines. The IC_50s_ in HT1080/ACE2 and A549/ACE2 cells for NYBSP-1 were 4.1±0.26 and 2.2±0.14, respectively, for NYBSP-2 were 2.9±0.27 and 2.68±0.14, and for NYBSP-4 1.97±0.14 and 2.8±0.08, respectively. NYBSP-3 was the least active peptide with an IC_50_ of 12.9±0.35 μM in HT1080/ACE cells and ∼25 μM in A549/ACE2 cells. Most significantly, as expected, neither of the two controls, the linear peptide NYBSP-C and the double-stapled peptide StRIP16 showed any antiviral activity while the IC_50_ detected for the anti-ACE2 mAb was >70 ng/mL (at this concentration, we detected about 65-80% viral inhibition). It is interesting to note that NYBSP-3 had the lowest % α-helicity among all the stapled peptide designed, and it also showed the least antiviral activity. Moreover, none of the peptides showed activity against the control pseudovirus VSV-G (**Table 1**), suggesting that their inhibitory activity is specific to SARS-CoV-2.

**Table 1.** Antiviral activity (IC_50_) of peptides in single cycle assay in HT1080/ACE2 and A549/ACE2 cells infected with pseudoviruses (NL4-3ΔEnv-NanoLuc/SARS-CoV-2 and NL4-3.Luc.R-E-/VSV-G) and cytotoxicity (CC_50_).

| Peptide | HT1080/ACE2 cells |  |  | A549/ACE2 cells |  |  |
| --- | --- | --- | --- | --- | --- | --- |
|  | IC<sub>50</sub> (μM)<sup>a</sup> |  | CC<sub>50</sub> (μM)<sup>a</sup> | IC<sub>50</sub> (μM)<sup>a</sup> |  | CC<sub>50</sub> (μM)<sup>a</sup> |
|  | SARS-CoV-2 | VSV-G |  | SARS-CoV-2 | VSV-G |  |
| NYBSP-1 | 4.1±0.26 | >27.5 | >27.5<sup>#</sup> | 2.2±0.14 | >27.5 | >27.5<sup>#</sup> |
| NYBSP-2 | 2.9±0.27 | >26.8 | >26.8<sup>#</sup> | 2.68±0.14 | >26.8 | >26.8<sup>#</sup> |
| NYBSP-3 | 12.9±0.35 | >27.6 | >27.6<sup>#</sup> | ~25 | >27.6 | >27.6<sup>#</sup> |
| NYBSP-4 | 1.97±0.14 | >26.6 | >26.6<sup>#</sup> | 2.8±0.08 | >26.6 | >26.6<sup>#</sup> |
| NYBSP-C* | >27.7 | >27.7 | >27.7<sup>#</sup> | >27.7 | >27.7 | >27.7<sup>#</sup> |
| StRIP16** | >43.5 | >43.5 | >43.5<sup>#</sup> | >43.5 | >43.5 | >43.5<sup>#</sup> |
| Anti-ACE2 | <70ng/ml | >5μg/ml | >5μg/ml<sup>#</sup> | <70ng/ml | >5μg/ml | >5μg/ml<sup>#</sup> |

<sup>a</sup> The reported IC<sub>50</sub> and CC<sub>50</sub> values represent the means ± standard deviations (n =3)
<sup>#</sup> <10% toxicity at this dose
\*NYBSP-C, a linear peptide from Helix-1 was used as a control
\*\* StRIP16, a ACE2-unrelated Rab8a GTPase-binding double-stapled peptide reported earlier <sup>29</sup>, was used as a control.

**Figure 7.**
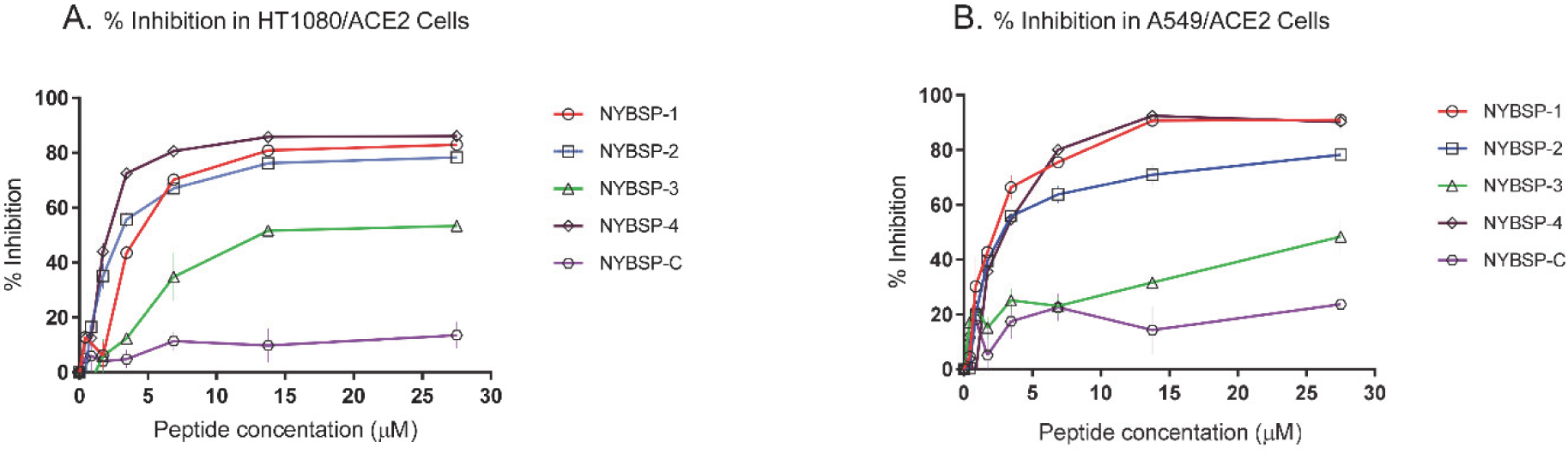
Dose-Response plots of the antiviral activity of peptides in a single-cycle assay performed in HT1080/ACE2 and A549/ACE2 cells infected with pseudovirus NL4-3ΔEnv-NanoLuc/SARS-CoV-2. Antiviral activity in (A) HT1080/ACE2 cells and (B) A549/ACE2 cells. Data on stapled peptides NYBSP-1, NYBSP-2, NYBSP-3, NYBSP-4, and a control Linear peptide NYBSP-C are shown as mean ± SD of three independent experiments.

The cytotoxicity (CC_50_) of the peptides was assessed in parallel with the inhibition assays. We found that the peptides did not induce any apparent cell toxicity at the highest dose tested. More specifically, <10% toxicity was detected at the higher dose used in this assay (**Table 1**).

Additionally, the antiviral activity of the peptides was evaluated by infecting Vero E6 cells with the replication-competent virus SARS-CoV-2 (US_WA-1/2020). The cells were observed under the microscope after 4 days of incubation to evaluate the formation of virus-induced cytopathic effect (CPE), and the efficacy of the peptides was expressed as the lowest concentration capable of completely preventing virus-induced CPE (IC_100_). We found that the linear peptide NYSBP-C and the double-stapled NYBSP-3 did not completely prevent the formation of the virus-induced CPE at the highest dose used in this assay (**Table 2**), while NYBSP-1 was the most efficient in preventing the complete formation of CPEs at an IC_100_ of 17.2 µM. NYBSP-2 and NYBSP-4 also prevented the formation of the virus-induced CPE with an IC_100_ of about 33 µM.

**Table 2.** Antiviral activity (IC_100_) of peptides in Vero E6 cells infected with SARS-CoV-2 (US_WA-1/2020).

| Peptide | IC<sub>100</sub> (μM)* |
| --- | --- |
| NYBSP-1 | 17.2 |
| NYBSP-2 | 33.5 |
| NYBSP-3 | NA** |
| NYBSP-4 | 33 |
| NYBSP-C | NA** |

\*Values indicate the lowest concentration capable of completely preventing virus-induced CPE in 100% of the wells.
\*\*NA means the presence of CPE at the highest concentration

## Methods

### Peptide synthesis

#### General procedure of coupling

Fmoc Rink Amide MBHA Resin (0.1 mmol) was swelled in DMF (20 mL) for 1 h. The suspension was filtered. 20% Piperidine in DMF (20 mL) was added to the resin to remove Fmoc. The suspension was kept at room temperature for 0.5 h, while a stream of nitrogen was bubbled through it. The mixture was filtered, and the resin was washed with DMF (6×20 mL). Fmoc-Asn(Trt)-OH (0.3 mmol) was pre-activated with DIC (0.3 mmol) and HOBt (0.3 mmol) in 10 mL DMF, then the mixture was added into the resin. The reaction was carried out in a nitrogen atmosphere. Kaiser ninhydrin test was used to indicate reaction completion. After the reaction was completed, the suspension was filtered, and the resin was washed with DMF (3×20 mL). After that, all sequences of amino acids were coupled and completed to give the peptidyl resin full protection, which was used to do the cyclization with Grubbs 1st catalyst.

The peptidyl resin was washed DCM (3×20mL). Then add 10 mL 1,2-dichloroethane in the reaction vessel, and the solution was bubbled by N_2_. 15 min later, 30 mg of Grubbs 1^st^ catalyst was added to the reaction vessel and bubbled by N_2_ for 4 h, filter, and the resin was washed with MeOH (2×150 mL), DCM (2×150 mL) and MeOH (2×150 mL). The resin was dried under vacuum overnight to give Cyclized peptidyl resin.

E solution (TFA:Thioanisole:phenol:EDT:H_2_O=87.5:5:2.5:2.5:2.5, 350 mL) was added into the peptidyl resin, and the suspension was shaken for 3.5 h followed by filtration. Ether (2000 mL) was added into the filtrate, and peptide precipitated. The mixture was centrifuged, and the ether layer was decanted. The peptide was washed three times with ether, dried under vacuum overnight to give the crude.

All crude peptides were purified by pre-HPLC to > 90% purity. They were acetylated at the C-terminus and amidated at the N-terminus. The molecular weight of all peptides was confirmed by electrospray mass spectrometry.

#### Circular Dichroism spectroscopy

The secondary structure analysis to calculate % α-helicity was performed by Creative-Biolabs (New York, NY, USA). In brief, peptides were dissolved in 10mM PBS to make a final concentration of 0.125 mg/ml in PBS. The CD spectra were obtained on a Chirascan spectropolarimeter (Applied Photophysics, Leatherhead, UK). The measurement parameters were set up as follows-Wavelength: (185-260nm); Step Size: 0.5nm; Temperature: 25 °C. The % helicity was calculated using the software CDNN (Circular Dichroism analysis using Neural Networks) (http://gerald-boehm.de/download/cdnn).^33^

#### SPR analysis

The binding study of one double-stapled peptide, NYBSP-4, and the linear control peptide, NYBSP-C, was performed by Profacgen, New York, NY. The bare gold-coated (thickness 47 nm) PlexArray Nanocapture Sensor Chip (Plexera Bioscience, Seattle, WA, US) was prewashed with 10× PBST for 10 min, 1× PBST for 10 min, and deionized water twice for 10 min before being dried under a stream of nitrogen prior to use. Various concentrations of biotinylated recombinant SARS-CoV-2 Spike RBD protein (Bioss Antibodies, Woburn, MA) dissolved in water were manually printed onto the Chip with Biodo bioprinting at 40% humidity via biotin-avidin conjugation. Each concentration was printed in replicate, and each spot contained 0.2 µL of the sample solution. The Chip was incubated in 80% humidity at 4°C overnight and rinsed with 10× PBST for 10 min, 1× PBST for 10 min, and deionized water twice for 10 min. The Chip was then blocked with 5% (w/v) non-fat milk in water overnight and washed with 10× PBST for 10 min, 1× PBST for 10 min, and deionized water twice for 10 min before being dried under a stream of nitrogen prior to use. SPRi measurements were performed with PlexAray HT (Plexera Bioscience, Seattle, WA, US). Collimated light (660 nm) passes through the coupling prism, reflects off the SPR-active gold surface, and is received by the CCD camera. Buffers and the test peptides were injected by a non-pulsatile piston pump into the 30 µL flowcell that was mounted on the coupling prim. Each measurement cycle contained four steps: washing with running buffer contained 10 mM HEPES, pH 7.4, 150 mM NaCl, 3 mM EDTA and 0.05% Tween-20 at a constant rate of 2 µL/s to obtain a stable baseline, sample injection at 5 µL/s for binding, surface washing with running buffer at 2 µL/s for 300 s, and regeneration with 0.5% (v/v) H3PO4 at 2 µL/s for 300 s. All the measurements were performed at 25°C. The signal changes after binding and washing (in AU) are recorded as the assay value.

#### Kinetics fitting and analysis

Selected protein-grafted regions in the SPR images were analyzed, and the average reflectivity variations of the chosen areas were plotted as a function of time. Real-time binding signals were recorded and analyzed by the Data Analysis Module (DAM, Plexera Bioscience, Seattle, WA, US). Kinetic analysis was performed using BIAevaluation 4.1 software (Biacore, Inc.). The resulting data were fit to a 1:1 binding model using Biacore Evaluation Software (GE Healthcare).

#### Plasma stability study

The plasma stability study was performed by Creative-Biolabs (New York, NY, USA). In brief, the pooled frozen human plasma (BioIVT, UK) was thawed in a water bath at 37 °C prior to the experiment. Plasma was centrifuged at 4000 rpm for 5 min, and the clots were removed, if any. The pH was adjusted to 7.4 ± 0.1 if required. 1 mM stock solution of the peptides was prepared with water with 5% NH_3_.H_2_O. 1 mM intermediate of positive control Propantheline was prepared by diluting 10 µL of the stock solution with 90 µL ultrapure water.

For control, 100 μM dosing solution was prepared by diluting 10 µL of the intermediate solution (1 mM) with 90 µL 45% MeOH/H_2_O. For test peptide, 100 μM dosing solution was prepared by diluting 10 µL of the intermediate solution (1 mM) with 90 µL DMSO. 98 µL of blank plasma was spiked with 2 μL of dosing solution (100 μM) to achieve 2 μM of the final concentration in duplicate, and samples were Incubated at 37 °C in a water bath. At each time point (0,10, 30, 60, and 120 min), add 100 μL 4% H_3_PO_4_ first, and then 800 μL of stop solution (200 ng/mL tolbutamide and 200 ng/mL Labetalol in MeOH) was added to precipitate protein and mixed thoroughly. Sample plates were centrifuged at 4000 rpm for 10 min. An aliquot of supernatant 100 μL was transferred from each well to the plate. The samples were shaken at 800 rpm for about 10 min before submitting to LC-MS/MS analysis. The % remaining of the test peptides after incubation in plasma was calculated using the following equation:

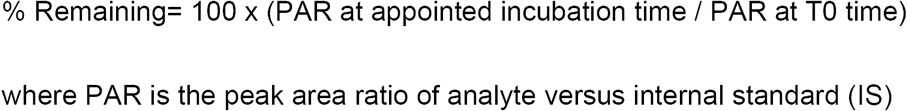

#### Cells and plasmids

The Human Lung Carcinoma Cells (A549) expressing Human Angiotensin-Converting Enzyme 2 (HA-FLAG) (Catalog No. NR-53522) were obtained from BEI Resources, NIAID, NIH. HeLa, A549, HT-1080, and HEK293T/17 cells were purchased from ATCC (<u>Manassas, VA</u>). The HT1080/ACE2 (human fibrosarcoma) cells and the two plasmids pNL4-3ΔEnv-NanoLuc and pSARS-CoV-2-S_Δ19_ were kindly provided by Dr. Paul Bieniasz of Rockefeller University^30^. The pHEF-VSV-G envelope expression vector ^34^ and the Env-deleted proviral backbone plasmids pNL4-3.Luc.R-E-DNA ^35, 36^ were obtained through the NIH ARP.

#### Pseudoviruses preparation

To prepare pseudoviruses capable of single-cycle infection, 10×10^6^ HEK293T/17 cells were transfected with a proviral backbone plasmid and an envelope expression vector by using FuGENE6 (Promega, <u>Madison, WI</u>) and following the manufacturer’s instructions. To obtain the SARS-CoV-2 pseudovirus, the cells were transfected with the HIV-1 Env-deleted proviral backbone plasmid pNL4-3ΔEnv-NanoLuc DNA and the pSARS-CoV-2-S_Δ19_ plasmid^30^. For the VSV-G pseudovirus, the cells were transfected with the Env-deleted proviral backbone plasmids pNL4-3.Luc.R-.E-DNA and the pHEF-VSV-G envelope expression vector. Pseudovirus-containing supernatants were collected two days after transfection, filtered, tittered, and stored in aliquots at −80 °C. Pseudovirus titers were determined to identify the 50% tissue culture infectious dose (TCID_50_) by infecting different cell types. For the titers in HT1080/ACE2 cells, 2×10^4^ cells were added to 100-μL aliquots of serial 2-fold dilutions of pseudoviruses in a 96-well plate and incubated for 24 h. For the titers in A549/ACE2 cells, 1×10^4^ cells were added to 100-μL aliquots of serial 2-fold dilutions of pseudoviruses in a 96-well plate and incubated for 48 h. Following the incubation time, the cells were washed with PBS and lysed with 50 μL of the cell culture lysis reagent (Promega, <u>Madison, WI</u>). For the SARS-CoV-2 titers, 25 µL of the lysates were transferred to a white plate and mixed with the same volume of Nano-Glo® Luciferase reagent (Promega). For the VSV-G titers, 25 µL of the lysates were transferred to a white plate and mixed with 50µL of luciferase assay reagent (Luciferase assay system, Promega). We immediately measured the luciferase activity with a Tecan SPARK multifunctional microplate reader (Tecan, Research Triangle Park, NC). The wells producing relative luminescence unit (RLU) levels 10 times the cell background were scored as positive. We calculated the TCID_50_ according to the Spearman-Karber method ^37^.

#### Analysis of the incorporation of spike proteins into SARS-CoV-2 pseudovirus

To confirm the incorporation of the spike protein into the SARS-CoV-2 pseudovirus, 2 mL of the pseudovirus-containing supernatant was ultra-centrifuged for 2 h at 38,000 rpm on a 20 % sucrose cushion to concentrate the viral particles. Viral pellets were lysed and processed for protein analysis. The viral proteins were resolved on a NuPAGE Novex 4–12 % Bis-Tris Gel (Invitrogen, <u>Carlsbad, CA</u>) and immuno-detected with a SARS Spike protein Antibody (NB-100-56578, Novus Biological, <u>Littleton, CO</u>). An anti-Rabbit-IgG HRP linked whole antibody (GE Healthcare, <u>Chicago, IL</u>) was used as a secondary antibody. Proteins were visualized using chemiluminescence. Additionally, the correlation of the pseudovirus SARS-CoV-2 with the expression of the ACE2 receptor was analyzed by infecting five different cell types expressing different amounts of the ACE2 receptor, with the same volume of the pseudovirus-containing supernatant. Briefly, 50 µL of SARS-CoV-2 diluted with 50 µL serum-free medium was added to wells of a 96-well cell culture plate. Next, the cells were added as follow: HT-1080\ACE2 and HT-1080 cells were added to the respective wells at the concentration of 2×10^4^ cells/well and incubated for 24 h at 37°C; A549\ACE2, A549, and Hela cells were added to the respective wells at a concentration of 1×10^4^ cells**/**well and incubated for 48 h at 37°C. Uninfected cells for all cell lines were used as a negative control. Following the incubation time, the cells were washed with PBS and lysed with 50 μL of the cell culture lysis reagent. 25 µL of the lysates were transferred to a white plate and mixed with the same volume of Nano-Glo® Luciferase reagent. The luciferase activity was immediately measured with a Tecan SPARK.

#### Evaluation of the ACE2 expression in different cell lines

The expression levels of the ACE2 receptor was evaluated by Western Blot to find a correlation with the infection levels detected in the five cell lines HT-1080\ACE2 and HT-1080, A549\ACE2, A549, and Hela. Cell pellets were lysed and processed for protein analysis. For Blot-1, we loaded 50 µg of proteins (Media, Hela, HT1080, and HT1080/ACE2), and for Blot-2, we loaded 75 µg of proteins (Media, Hela, A549 and A549/ACE2). The proteins were resolved on a NuPAGE Novex 4–12 % Bis-Tris Gel and immuno-detected with a human anti-ACE2 mAb (AC384) (Adipogen Life Sciences, San Diego, CA). The ECL Mouse IgG, HRP-linked whole Ab (from sheep) (Amersham, <u>Little Chalfont, UK</u>) was used as a secondary antibody. Proteins were also immuno-detected with the housekeeping gene β-actin as a loading control. Proteins were visualized using chemiluminescence.

## Measurement of antiviral activity

### HT1080/ACE2 cells

The antiviral activity of the peptides was evaluated in single-cycle infection assay by infecting HT1080/ACE2 cells with the SARS-CoV-2 pseudovirus, as previously described with minor modifications ^31^. Briefly, in 96-well culture plates, aliquots of SARS-CoV-2 at about 3000 TCID_50_/well at a multiplicity of infection (MOI) of 0.1, were pre-incubated with escalating concentrations of the peptides for 30 min. Next, 2×10^4^ cells were added to each well and incubated at 37°C. HT1080/ACE2 cells cultured with medium with or without the SARS-CoV-2 pseudovirus were included as positive and negative controls, respectively. The linear peptide NYBSP-C, StRIP16 ^29^ (Tocris, <u>Bristol, UK</u>) which is a SARS-CoV-2-unrelated Rab8a GTPase-binding double-stapled peptide and an anti-ACE2 mAb (AC384; Catalog# AG-20A-0037PF, Adipogen Life Sciences, San Diego, CA) were used as controls. In the case of the anti-ACE2 mAb, the cells were pre-incubated with escalating concentrations of the mAb for 30 min, then infected with the pseudovirus SARS-CoV-2. Following 24 h incubation, the cells were washed with 100 µL of PBS and lysed with 50 µL of lysis buffer (Promega). 25 µL of the lysates were transferred to a white plate and mixed with the same volume of Nano-Glo® Luciferase reagent (Promega). The luciferase activity was measured immediately with the Tecan SPARK. The percent inhibition by the peptides and the IC_50_ (the half-maximal inhibitory concentration) values were calculated using the GraphPad Prism 7.0 software (San Diego, CA). Additionally, to test the specificity of the double-stapled peptides, we evaluated their activity against pseudovirus VSV-G by following the infection protocol described above. Following 24 h incubation, 25 µL of the lysates were transferred to a white plate and mixed with 50 µL of a luciferase assay reagent. The luciferase activity was immediately measured.

### A549/ACE2 cells

For the evaluation of the antiviral activity in A549/ACE2 cells, aliquots of the pseudovirus SARS-CoV-2 at about 1500 TCID_50_/well at a MOI of 0.1, were pre-incubated with escalating concentrations of the double-stapled peptides for 30 min. Next, 1×10^4^ cells were added to each well and incubated. A549/ACE2 cells cultured with medium with or without the SARS-CoV-2 pseudovirus were included as positive and negative controls, respectively. The linear peptide NYBSP-C, the double-stapled peptide StRIP16, and the anti-ACE2 mAb were used as controls. In the case of the anti-ACE2 mAb, the cells were pre-incubated with escalating concentrations of the Ab for 30 min, then infected with the pseudovirus SARS-CoV-2. Following 48 h incubation, the cells were washed with PBS and lysed with 50 µL of lysis buffer. 25 µL of the cell lysates were processed as reported above to measure the luciferase activity and calculate the percent inhibition by the peptides and the IC_50_. Moreover, the double-stapled peptides were evaluated against pseudovirus VSV-G by following the infection protocol described above. Following 48 h incubation, 25 µL of the lysates were transferred to a white plate and mixed with 50 µL of a luciferase assay reagent. The luciferase activity was immediately measured.

### SARS-CoV-2 Microneutralization Assay

The standard live virus-based microneutralization (MN) assay was used ^38-40^. Briefly, serially two-fold and duplicate dilutions of individual peptides were incubated with 120 plaque-forming unit (PFU) of SARS-CoV-2 (US_WA-1/2020) at room temperature for 1 h before transferring into designated wells of confluent Vero E6 cells (ATCC, CRL-1586) grown in 96-well cell culture plates. Vero E6 cells cultured with medium with or without the same amount of virus were included as positive and negative controls, respectively. After incubation at 37°C for 3-4 days, individual wells were observed under the microscope to determine the virus-induced formation of cytopathic effect (CPE). The efficacy of individual drugs was expressed as the lowest concentration capable of completely preventing virus-induced CPE in 100% of the wells.

### Evaluation of cytotoxicity

#### HT1080/ACE2 cells

The evaluation of the cytotoxicity of the peptides in HT1080/ACE2 cells was performed in parallel with the antiviral activity assay and measured by using the colorimetric CellTiter 96® AQueous One Solution Cell Proliferation Assay (MTS) (Promega, <u>Madison, WI</u>) following the manufacturer’s instructions. Briefly, aliquots of 100 µL of the peptides at graded concentrations were incubated with 2 x 10^4^ / well HT1080/ACE2 cells and cultured at 37 °C. Following 24 h incubation, the MTS reagent was added to the cells and incubated for 4 h at 37 °C. The absorbance was recorded at 490 nm. The percent of cytotoxicity and the CC_50_ (the concentration for 50 % cytotoxicity) values were calculated as above.

#### A549/ACE2 cells

For the cytotoxicity assay in A549/ACE2 cells, aliquots of escalating concentrations of the peptides were incubated with 1 x 10^4^ / well A549/ACE2 cells and cultured at 37 °C. Following 48 h incubation, the MTS reagent was added to the cells and incubated for 4 h at 37 °C. The absorbance was recorded at 490 nm. The percent of cytotoxicity and the CC_50_ values were calculated as above.

## Conclusions

The current pandemic made it urgent to develop therapeutics urgently against COVID-19. We took advantage of the structural knowledge of the binding site of SARS-CoV-2 RBD and the ACE2 receptor. We envisioned that stapled peptide based on the helix region of ACE2 that binds with the SARS-CoV-2 RBD might work as a decoy receptor and bind the SARS-CoV-2 preferentially and prevent its entry to the host cells and thereby, the subsequent infection of the virus. We reported the successful design of double-stapled peptides with potent antiviral activity in HT1080/ACE2 and human lung carcinoma cells, A549/ACE2. Most significantly, all three active stapled peptides with potent antiviral activity against SARS-CoV-2 pseudovirus showed high helical content (60-94%). We also evaluated the antiviral activity of the peptides by infecting Vero E6 cells with the replication-competent authentic SARS-CoV-2 (US_WA-1/2020). NYBSP-1, NYBSP-2, and NYBSP-4 completely (IC_100_) inhibited virus-induced CPE at low µM doses. One of the most active stapled peptides, NYBSP-4, demonstrated a binding affinity (K_D_) of 2.2 µM with SARS-CoV-2 RBD, which was in line with the antiviral activity of the stapled peptide against SARS-CoV-2 pseudovirus. NYBSP-4 also showed substantial resistance to degradation by proteolytic enzymes in human plasma. The lead stapled peptides are expected to pave the way for further optimization of a clinical candidate.

## Acknowledgment

The study was conducted using an intramural fund to AKD from the New York Blood Center. We gratefully acknowledge the generous gift of cells (highly expressed ACE2) and several plasmids to create the pseudovirus from Dr. Paul Bieniasz of the Rockefeller University.

## References

1. Batlle, D.; Wysocki, J., Satchell, K., Soluble angiotensin-converting enzyme 2: a potential approach for coronavirus infection therapy? Clin Sci (Lond) 2020, 134 (5), 543–545.

2. Recombinant human angiotensin-converting enzyme 2 (rhACE2) as a treat-ment for patients with COVID-19. https://www.clinicaltrialsregister.eu/ctr-search/search?query=eudract_number:2020-001172-15.

3. Wysocki, J., Ye, M., Rodriguez, E., Gonzalez-Pacheco, F. R., Barrios, C., Evora, K., Schuster, M., Loibner, H., Brosnihan, K. B., Ferrario, C. M., Penninger, J. M., Batlle, D., Targeting the degradation of angiotensin II with recombinant angiotensin-converting enzyme 2: prevention of angiotensin II-dependent hypertension. Hypertension 2010, 55 (1), 90–8.

4. Haschke, M., Schuster, M., Poglitsch, M., Loibner, H., Salzberg, M., Bruggisser, M., Penninger, J., Krahenbuhl, S., Pharmacokinetics and pharmacodynamics of recombinant human angiotensin-converting enzyme 2 in healthy human subjects. Clin Pharmacokinet 2013, 52 (9), 783–92.

5. Lei, C., Qian, K., Li, T., Zhang, S., Fu, W., Ding, M., Hu, S., Neutralization of SARS-CoV-2 spike pseudotyped virus by recombinant ACE2-Ig. Nat Commun 2020, 11 (1), 2070.

6. Zhang, G., Pomplun, S., Loftis, A. R., Tan, X., Loas, A., Pentelute, B. L., The first-in-class peptide binder to the SARS-CoV-2 spike protein. BioRxiv 2020.

7. Walls, A. C., Park, Y. J., Tortorici, M. A., Wall, A., McGuire, A. T., Veesler, D., Structure, Function, and Antigenicity of the SARS-CoV-2 Spike Glycoprotein. Cell 2020, 181 (2), 281–292 e6.

8. Wrapp, D., Wang, N., Corbett, K. S., Goldsmith, J. A., Hsieh, C. L., Abiona, O., Graham, B. S., McLellan, J. S., Cryo-EM structure of the 2019-nCoV spike in the prefusion conformation. Science 2020, 367 (6483), 1260–1263.

9. Lan, J., Ge, J., Yu, J., Shan, S., Zhou, H., Fan, S., Zhang, Q., Shi, X., Wang, Q., Zhang, L., Wang, X., Structure of the SARS-CoV-2 spike receptor-binding domain bound to the ACE2 receptor. Nature 2020, 581 (7807), 215–220.

10. Shang, J., Ye, G., Shi, K., Wan, Y., Luo, C., Aihara, H., Geng, Q., Auerbach, A., Li, F., Structural basis of receptor recognition by SARS-CoV-2. Nature 2020, 581 (7807), 221–224.

11. Ali, A. M., Atmaj, J., Van Oosterwijk, N., Groves, M. R., Domling, A., Stapled Peptides Inhibitors: A New Window for Target Drug Discovery. Comput Struct Biotechnol J 2019, 17, 263–281.

12. Iegre, J., Ahmed, N. S., Gaynord, J. S., Wu, Y., Herlihy, K. M., Tan, Y. S., Lopes-Pires, M. E., Jha, R., Lau, Y. H., Sore, H. F., Verma, C., DH, O. D., Pugh, N., Spring, D. R., Stapled peptides as a new technology to investigate protein-protein interactions in human platelets. Chem Sci 2018, 9 (20), 4638–4643.

13. Moiola, M., Memeo, M. G., Quadrelli, P., Stapled Peptides-A Useful Improvement for Peptide-Based Drugs. Molecules 2019, 24 (20).

14. Muppidi, A., Zhang, H., Curreli, F., Li, N., Debnath, A. K., Lin, Q., Design of antiviral stapled peptides containing a biphenyl cross-linker. Bioorg Med Chem Lett 2014, 24 (7), 1748–51.

15. Charoenpattarapreeda, J., Tan, Y. S., Iegre, J., Walsh, S. J., Fowler, E., Eapen, R. S., Wu, Y., Sore, H. F., Verma, C. S., Itzhaki, L., Spring, D. R., Targeted covalent inhibitors of MDM2 using electrophile-bearing stapled peptides. Chem Commun (Camb) 2019, 55 (55), 7914–7917.

16. Cowell, J. K., Teng, Y., Bendzunas, N. G., Ara, R., Arbab, A. S., Kennedy, E. J., Suppression of Breast Cancer Metastasis Using Stapled Peptides Targeting the WASF Regulatory Complex. Cancer Growth Metastasis 2017, 10, 1179064417713197.

17. Cromm, P. M., Spiegel, J., Grossmann, T. N., Hydrocarbon stapled peptides as modulators of biological function. ACS Chem Biol 2015, 10 (6), 1362–75.

18. Dietrich, L., Rathmer, B., Ewan, K., Bange, T., Heinrichs, S., Dale, T. C., Schade, D., Grossmann, T. N., Cell Permeable Stapled Peptide Inhibitor of Wnt Signaling that Targets beta-Catenin Protein-Protein Interactions. Cell Chem Biol 2017, 24 (8), 958–968 e5.

19. Dougherty, P. G., Wen, J., Pan, X., Koley, A., Ren, J. G., Sahni, A., Basu, R., Salim, H., Appiah Kubi, G., Qian, Z., Pei, D., Enhancing the Cell Permeability of Stapled Peptides with a Cyclic Cell-Penetrating Peptide. J Med Chem 2019, 62 (22), 10098–10107.

20. Gaillard, V., Galloux, M., Garcin, D., Eleouet, J. F., Le Goffic, R., Larcher, T., Rameix-Welti, M. A., Boukadiri, A., Heritier, J., Segura, J. M., Baechler, E., Arrell, M., Mottet-Osman, G., Nyanguile, O., A Short Double-Stapled Peptide Inhibits Respiratory Syncytial Virus Entry and Spreading. Antimicrob Agents Chemother 2017, 61 (4).

21. Verdine, G. L., Hilinski, G. J., Stapled peptides for intracellular drug targets. Methods Enzymol 2012, 503, 3–33.

22. Clinical Programs: ALRN-6924. https://www.aileronrx.com/clinical-programs/.

23. Mourtada, R., Herce, H. D., Yin, D. J., Moroco, J. A., Wales, T. E., Engen, J. R., Walensky, L. D., Design of stapled antimicrobial peptides that are stable, nontoxic and kill antibiotic-resistant bacteria in mice. Nat Biotechnol 2019, 37 (10), 1186–1197.

24. Ran, X., Liu, L., Yang, C. Y., Lu, J., Chen, Y., Lei, M., Wang, S., Design of High-Affinity Stapled Peptides To Target the Repressor Activator Protein 1 (RAP1)/Telomeric Repeat-Binding Factor 2 (TRF2) Protein-Protein Interaction in the Shelterin Complex. J Med Chem 2016, 59 (1), 328–34.

25. Bird, G. H., Crannell, W. C., Walensky, L. D., Chemical synthesis of hydrocarbon-stapled peptides for protein interaction research and therapeutic targeting. Curr Protoc Chem Biol 2011, 3 (3), 99–117.

26. Walensky, L. D., Kung, A. L., Escher, I., Malia, T. J., Barbuto, S., Wright, R. D., Wagner, G., Verdine, G. L., Korsmeyer, S. J., Activation of apoptosis in vivo by a hydrocarbon-stapled BH3 helix. Science 2004, 305 (5689), 1466–70.

27. Walensky, L. D., Bird, G. H., Hydrocarbon-stapled peptides: principles, practice, and progress. J Med Chem 2014, 57 (15), 6275–88.

28. Bird, G. H., Madani, N., Perry, A. F., Princiotto, A. M., Supko, J. G., He, X., Gavathiotis, E., Sodroski, J. G., Walensky, L. D., Hydrocarbon double-stapling remedies the proteolytic instability of a lengthy peptide therapeutic. Proc Natl Acad Sci U S A 2010, 107 (32), 14093–8.

29. Cromm, P. M., Spiegel, J., Kuchler, P., Dietrich, L., Kriegesmann, J., Wendt, M., Goody, R. S., Waldmann, H., Grossmann, T. N., Protease-Resistant and Cell-Permeable Double-Stapled Peptides Targeting the Rab8a GTPase. ACS Chem Biol 2016, 11 (8), 2375–82.

30. Schmidt, F., Weisblum, Y., Muecksch, F., Hoffmann, H. H., Michailidis, E., Lorenzi, J. C. C., Mendoza, P., Rutkowska, M., Bednarski, E., Gaebler, C., Agudelo, M., Cho, A., Wang, Z., Gazumyan, A., Cipolla, M., Caskey, M., Robbiani, D. F., Nussenzweig, M. C., Rice, C. M., Hatziioannou, T., Bieniasz, P. D., Measuring SARS-CoV-2 neutralizing antibody activity using pseudotyped and chimeric viruses. J Exp Med 2020, 217 (11).

31. Nie, J., Li, Q., Wu, J., Zhao, C., Hao, H., Liu, H., Zhang, L., Nie, L., Qin, H., Wang, M., Lu, Q., Li, X., Sun, Q., Liu, J., Fan, C., Huang, W., Xu, M., Wang, Y., Establishment and validation of a pseudovirus neutralization assay for SARS-CoV-2. Emerg Microbes Infect 2020, 9 (1), 680–686.

32. Smith, T. R. F., Patel, A., Ramos, S., Elwood, D., Zhu, X., Yan, J., Gary, E. N., Walker, S. N., Schultheis, K., Purwar, M., Xu, Z., Walters, J., Bhojnagarwala, P., Yang, M., Chokkalingam, N., Pezzoli, P., Parzych, E., Reuschel, E. L., Doan, A., Tursi, N., Vasquez, M., Choi, J., Tello-Ruiz, E., Maricic, I., Bah, M. A., Wu, Y., Amante, D., Park, D. H., Dia, Y., Ali, A. R., Zaidi, F. I., Generotti, A., Kim, K. Y., Herring, T. A., Reeder, S., Andrade, V. M., Buttigieg, K., Zhao, G., Wu, J. M., Li, D., Bao, L., Liu, J., Deng, W., Qin, C., Brown, A. S., Khoshnejad, M., Wang, N., Chu, J., Wrapp, D., McLellan, J. S., Muthumani, K., Wang, B., Carroll, M. W., Kim, J. J., Boyer, J., Kulp, D. W., Humeau, L., Weiner, D. B., Broderick, K. E., Immunogenicity of a DNA vaccine candidate for COVID-19. Nat Commun 2020, 11 (1), 2601.

33. Bohm, G., Muhr, R., Jaenicke, R., Quantitative analysis of protein far UV circular dichroism spectra by neural networks. Protein Eng 1992, 5 (3), 191–5.

34. Chang, L. J., Urlacher, V., Iwakuma, T., Cui, Y., Zucali, J., Efficacy and safety analyses of a recombinant human immunodeficiency virus type 1 derived vector system. Gene Ther 1999, 6 (5), 715–28.

35. Connor, R. I., Chen, B. K., Choe, S., Landau, N. R., Vpr is required for efficient replication of human immunodeficiency virus type-1 in mononuclear phagocytes. Virol 1995, 206 (2), 935–944.

36. He, J., Choe, S., Walker, R., di, M. P., Morgan, D. O., Landau, N. R., Human immunodeficiency virus type 1 viral protein R (Vpr) arrests cells in the G2 phase of the cell cycle by inhibiting p34cdc2 activity. J.Virol. 1995, 69 (11), 6705–6711.

37. https://www.hanc.info/labs/labresources/procedures/ACTGIMPAACT%20Lab%20Manual%20Archive/TCID50%20Determination%20-%20ARCHIVED_WM.pdf.

38. Li, W., Schafer, A., Kulkarni, S. S., Liu, X., Martinez, D. R., Chen, C., Sun, Z., Leist, S. R., Drelich, A., Zhang, L., Ura, M. L., Berezuk, A., Chittori, S., Leopold, K., Mannar, D., Srivastava, S. S., Zhu, X., Peterson, E. C., Tseng, C. T., Mellors, J. W., Falzarano, D., Subramaniam, S., Baric, R. S., Dimitrov, D. S., High Potency of a Bivalent Human VH Domain in SARS-CoV-2 Animal Models. Cell 2020, 183 (2), 429–441 e16.

39. Du, L., Zhao, G., Yang, Y., Qiu, H., Wang, L., Kou, Z., Tao, X., Yu, H., Sun, S., Tseng, C. T., Jiang, S., Li, F., Zhou, Y., A conformation-dependent neutralizing monoclonal antibody specifically targeting receptor-binding domain in Middle East respiratory syndrome coronavirus spike protein. J Virol 2014, 88 (12), 7045–53.

40. Agrawal, A. S., Ying, T., Tao, X., Garron, T., Algaissi, A., Wang, Y., Wang, L., Peng, B. H., Jiang, S., Dimitrov, D. S., Tseng, C. T., Passive Transfer of A Germline-like Neutralizing Human Monoclonal Antibody Protects Transgenic Mice Against Lethal Middle East Respiratory Syndrome Coronavirus Infection. Sci Rep 2016, 6, 31629.

